# Imaging-Based Age- and Sex-Related Cardiovascular and Skeletal Phenotypes in *Fbn1^C1041G/+^* Mice

**DOI:** 10.64898/2026.07.21.739835

**Authors:** Wenxin Liu, Shuolei Li, Xiangyu Zhang, Lina Su, Hang Lin, Peida Li, Jingying Liu, Yidong Niu, Sufang Li

**Author notes:** These authors contributed equally to this work (Wenxin Liu, Shuolei Li, Xiangyu Zhang). Corresponding authors: Sufang Li,; Yidong Niu,.

## Abstract

Marfan syndrome is a multisystem connective tissue disorder with age- and sex-related phenotypic variability. This study characterized early cardiovascular, valvular, transmitral filling, and skeletal phenotypes in *Fbn1^C1041G/+^* mice. Male and female wild-type and *Fbn1^C1041G/+^* mice were examined in independent age-specific cohorts from 4 to 20 weeks of age. Transthoracic echocardiography was used to measure aortic root and ascending aortic diameters, anterior mitral leaflet length, and mitral inflow Doppler parameters. Lateral X-ray imaging was used to quantify kyphosis angle. *Fbn1^C1041G/+^* mice of both sexes showed age-related aortic enlargement, with segment-specific sex-related heterogeneity involving the annulus and sinus. In mutant mice, anterior mitral leaflet elongation was detected from 8 weeks in males and at 20 weeks in females. Reduced E-wave velocity and E/A ratio were observed in both sexes at 4 weeks, with E-wave velocity remaining lower at most subsequent ages and a lower E/A ratio present at all ages; A-wave velocity did not differ between genotypes at any examined age. Reduced kyphosis angle was detected from 4 weeks in females and from 12 weeks in males. These findings show that early Fbn1-related phenotypes follow organ-specific age-related patterns, supporting age-resolved and sex-stratified assessment in preclinical Marfan syndrome studies.

## Introduction

Marfan syndrome (MFS) is an autosomal dominant connective tissue disorder caused by heterozygous mutations in *FBN1*, which encodes fibrillin-1, a major extracellular matrix (ECM) glycoprotein and an essential component of elastic microfibrils [1]. Pathogenic *FBN1* mutations impair microfibril assembly and function, compromising connective tissue integrity [2–4]. The major clinical manifestations involve the cardiovascular, skeletal, and ocular systems, with key disease features including progressive aortic dilation, valvular abnormalities, and skeletal deformities [5].

The *Fbn1^C1041G/+^* mouse, carrying a heterozygous p.C1041G mutation in Fbn1, is a well-established model of MFS that recapitulates major cardiovascular manifestations, including progressive aortic dilation and mitral valve disease [4,6]. Previous studies have shown that thoracic aortopathy in this model varies with age, aortic segment, and sex. Male mice generally exhibit more severe remodeling, whereas females show relative protection or delayed dilation [7,8]. Similar age- and sex-related variability has been reported in human MFS cohorts [9]. The early segment-specific age-related patterns of aortic dilation in male and female *Fbn1^C1041G/+^*mice, however, have not been fully defined. Characterizing these patterns may improve the interpretation of early disease manifestations and help identify age windows suitable for mechanistic or interventional studies.

Aortic root aneurysm and dissection represent the most life-threatening cardiovascular manifestations of MFS [10], but myxomatous thickening and elongation of the mitral valve leaflets are common and may impair cardiac function, especially when accompanied by clinically significant regurgitation [11,12]. Recent studies in *Fbn1^C1041G/+^* mice have demonstrated progressive alterations in mitral valve ECM composition, tissue mechanics, and function [13]. Mitral inflow Doppler parameters, including peak early diastolic filling velocity (E wave), peak late diastolic filling velocity during atrial contraction (A wave), and the E/A ratio, are routinely used to assess left ventricular diastolic filling [14]. Altered inflow profiles have been reported in MFS patients and in mouse models, suggesting abnormal transmitral filling patterns [13,15]. Whether mitral leaflet elongation and transmitral filling abnormalities are evident at similar ages during early disease and whether their age-related patterns differ between sexes remain unclear.

Skeletal manifestations, particularly kyphosis and scoliosis, are common features of MFS and contribute substantially to morbidity [16,17]. Previous studies in Marfan mouse models, including *Fbn1^C1041G/+^*mice, have described bone morphology, tissue material properties, and hyperkyphotic deformity [18–20]. The early postnatal age-related pattern of kyphosis and its possible sex dependence have received less attention.

Non-invasive imaging is essential for clinical assessment and longitudinal surveillance in MFS and offers a practical approach for phenotyping preclinical models. Nevertheless, early multi-system phenotyping from 4 weeks of age remains incompletely characterized in *Fbn1^C1041G/+^*mice, particularly in an age- and sex-stratified manner. To address this, we performed imaging-based phenotypic profiling of these mice from 4 to 20 weeks of age. By systematically measuring segment-specific aortic dimensions, AML length, mitral inflow Doppler parameters, and kyphosis angle, we aimed to define the age-related patterns of cardiovascular and skeletal phenotypes and to assess whether these patterns differ between sexes.

## Results

### General physiological parameters

Body weight and heart rate were comparable between WT and *Fbn1^C1041G/+^* mice in either sex from 4 to 20 weeks of age (Supplementary Fig. 1).

### Age- and sex-related differences in aortic root and ascending aortic dimensions in *Fbn1^C1041G/+^* mice

To assess age-related aortic enlargement and potential sex-related heterogeneity, we measured diameters of the aortic annulus, sinus, STJ and ascending aorta by transthoracic echocardiography from 4 to 20 weeks of age (Fig. 1a). In both genotypes and sexes, all four aortic segments showed significant enlargement with age (Fig. 1b-e and Table 1).

**Fig 1.**
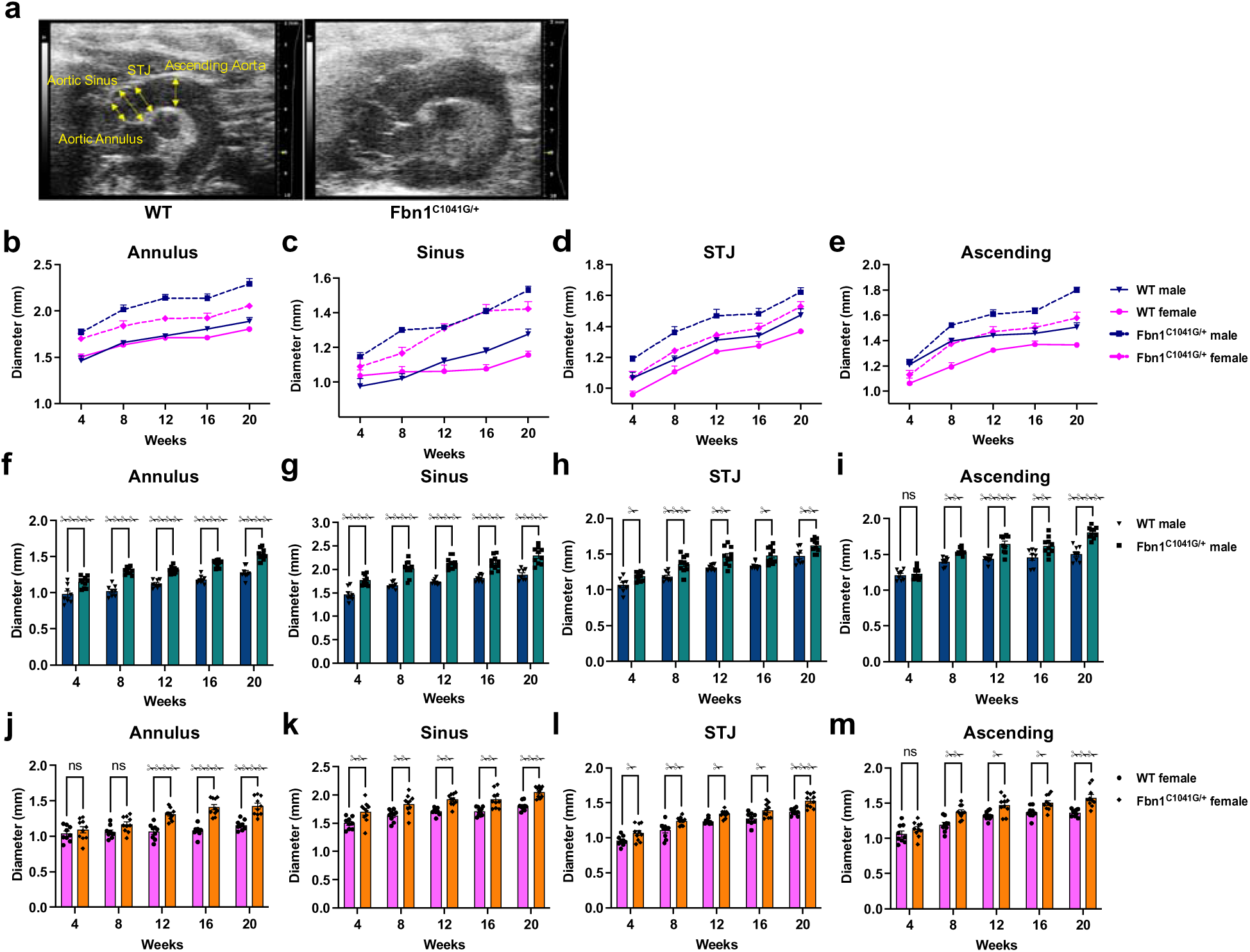
Age-related aortic root and ascending aortic dilation in male and female *Fbn1^C1041G/+^* mice. (a) Representative transthoracic echocardiographic images of the aortic root and ascending aorta in WT and *Fbn1^C1041G/+^* mice at 20 weeks of age. (b-e) Line charts showing age-related changes in aortic root and ascending aortic diameters in male and female WT and *Fbn1^C1041G/+^*mice. (f-m) Quantification of aortic annulus (f and j), aortic sinus (g and k), STJ (h and l), and ascending aortic (i and m) diameters in male (f-i) and female (j-m) WT and *Fbn1^C1041G/+^* mice at 4, 8, 12, 16, and 20 weeks of age. Data are presented as mean ± SEM (n = 8 WT male, 9 WT female, 10 *Fbn1^C1041G/+^*male, and 10 *Fbn1^C1041G/+^* female per age group). Statistical significance was determined by two-way ANOVA followed by Sidak’s multiple comparisons test. *P < 0.05, **P < 0.01, ***P < 0.001, ****P < 0.0001 versus age-matched WT mice; ns, not significant; STJ, sinotubular junction.

**Table 1.** Linear trend analysis of aortic dimensions by genotype and sex in mice aged 4 to 20 weeks.

| Aortic segment | Group | Slope, mm/week | 95% CI | P for trend |
| --- | --- | --- | --- | --- |
| Aortic Annulus | WT male | 0.0189 | 0.0143 to 0.0235 | <0.0001 |
|  | WT female | 0.0064 | 0.0016 to 0.0111 | 0.0095 |
|  | *Fbn1*<sup>C1041G/+</sup> male | 0.0219 | 0.0186 to 0.0252 | <0.0001 |
|  | *Fbn1*<sup>C1041G/+</sup> female | 0.0227 | 0.0167 to 0.0287 | <0.0001 |
| Aortic Sinus | WT male | 0.0251 | 0.0194 to 0.0307 | <0.0001 |
|  | WT female | 0.0169 | 0.0122 to 0.0217 | <0.0001 |
|  | *Fbn1*<sup>C1041G/+</sup> male | 0.0292 | 0.0215 to 0.0369 | <0.0001 |
|  | *Fbn1*<sup>C1041G/+</sup> female | 0.0198 | 0.0126 to 0.0270 | <0.0001 |
| Sinotubular Junction | WT male | 0.0242 | 0.0197 to 0.0287 | <0.0001 |
|  | WT female | 0.0246 | 0.0204 to 0.0288 | <0.0001 |
|  | *Fbn1*<sup>C1041G/+</sup> male | 0.0246 | 0.0192 to 0.0301 | <0.0001 |
|  | *Fbn1*<sup>C1041G/+</sup> female | 0.0265 | 0.0220 to 0.0311 | <0.0001 |
| Ascending Aorta | WT male | 0.0163 | 0.0110 to 0.0216 | <0.0001 |
|  | WT female | 0.0196 | 0.0144 to 0.0247 | <0.0001 |
|  | *Fbn1*<sup>C1041G/+</sup> male | 0.0312 | 0.0260 to 0.0365 | <0.0001 |
|  | *Fbn1*<sup>C1041G/+</sup> female | 0.0256 | 0.0193 to 0.0319 | <0.0001 |

**Note.** Values represent group-specific cross-sectional linear trend analyses of aortic dimensions from 4 to 20 weeks of age. The slope represents the estimated difference in aortic diameter associated with a one-week increase in age, with the 95% confidence interval (CI) shown. P for trend was obtained from simple linear regression using age as a continuous variable. Aortic dimensions and slopes are expressed in millimeters and millimeters per week of age, respectively. WT male, n = 8; WT female, n = 9; *Fbn1*<sup>C1041G/+</sup> male, n = 10; *Fbn1*<sup>C1041G/+</sup> female, n = 10 per age group.

Three-way ANOVA of aortic dimensions revealed segment-specific sex-related interactions. For the aortic annulus, the genotype × sex × age interaction was significant (P = 0.0062; Supplementary Table 1), indicating that the sex difference in the genotype-associated annular effect varied across age groups. For the sinus, the genotype × sex interaction was significant (P = 0.0001), whereas the genotype × sex × age interaction was not (P = 0.9388; Supplementary Table 1), indicating an overall sex difference in the genotype-associated sinus effect without evidence that this difference varied across age groups. For the STJ and ascending aorta, neither the genotype × sex nor the genotype × sex × age interaction was significant (both P > 0.05; Supplementary Table 1). As a focused secondary analysis, we also calculated the WT-corrected 4-to-20-week age-group contrasts. The largest corrected contrasts were observed in the male ascending aorta (0.27□mm) and the female annulus (0.21□mm); however, male-female differences in WT-corrected contrasts were not significant for any aortic segment (Supplementary Table 2).

At individual age groups, male *Fbn1^C1041G/+^* mice had larger diameters than WT in all root segments at each examined age, whereas the ascending aortic diameter was significantly larger than WT at 8, 12, 16, and 20 weeks (Fig. 1f–i). In females, sinus and STJ diameters were consistently larger than those of WT mice across all examined age groups, while significant genotype differences in the ascending aorta were observed at 8, 12, 16, and 20 weeks and those in the annulus at 12, 16, and 20 weeks (Fig. 1j–m).

To assess the potential confounding effect of individual body size variation on aortic dimensions, we further performed multiple linear regression analyses with body weight as a covariate within each age-sex subgroup (Supplementary Table 3). The results were largely consistent with the unadjusted analyses across all subgroups, except for 8-week-old females, in which a significant genotype effect emerged after weight adjustment in the aortic annulus, whereas the unadjusted comparison was not significant. This result suggests that inter-individual body weight variability partially masked the genotype effect in this specific subgroup. Overall, the weight-adjusted analyses were highly consistent with our primary findings, suggesting that the observed aortic differences were not substantially explained by individual body weight.

We also examined physiological sex differences in WT aortas. The aortic annulus was significantly larger in males than in females at 20 weeks, with a similar but non-significant difference at 16 weeks (Supplementary Fig. 2a). Sinus diameters showed no sex differences at any age (Supplementary Fig. 2b). STJ diameters were larger in males at 4 and 20 weeks, whereas differences at 8-16 weeks did not reach statistical significance (Supplementary Fig. 2c). The most consistent sex difference was in the ascending aorta, where males had significantly larger diameters at 4, 8, 12 and 20 weeks (Supplementary Fig. 2d). These physiological sex differences suggest that absolute aortic diameters in mutant mice should be interpreted cautiously in direct cross-sex comparisons.

### Age- and sex-related differences in AML length in *Fbn1^C1041G/+^* mice

We measured anterior mitral leaflet (AML) length by transthoracic echocardiography at 4, 8, 12, 16, and 20 weeks of age (Fig. 2a). AML length increased with age in both genotypes and sexes, and the cross-sectional slopes in mutants were approximately twice those in WT controls (Fig. 2b and Table 2). Three-way ANOVA showed significant main effects of genotype and age (both P < 0.0001), whereas neither the genotype × sex interaction (P = 0.3005) nor the genotype × sex × age interaction (P = 0.6744) reached significance (Supplementary Table 4). These results indicate that the *Fbn1^C1041G/+^* mutation drives a robust leaflet elongation across ages, without evidence that this age-related genotypic effect differs between males and females.

**Fig 2.**
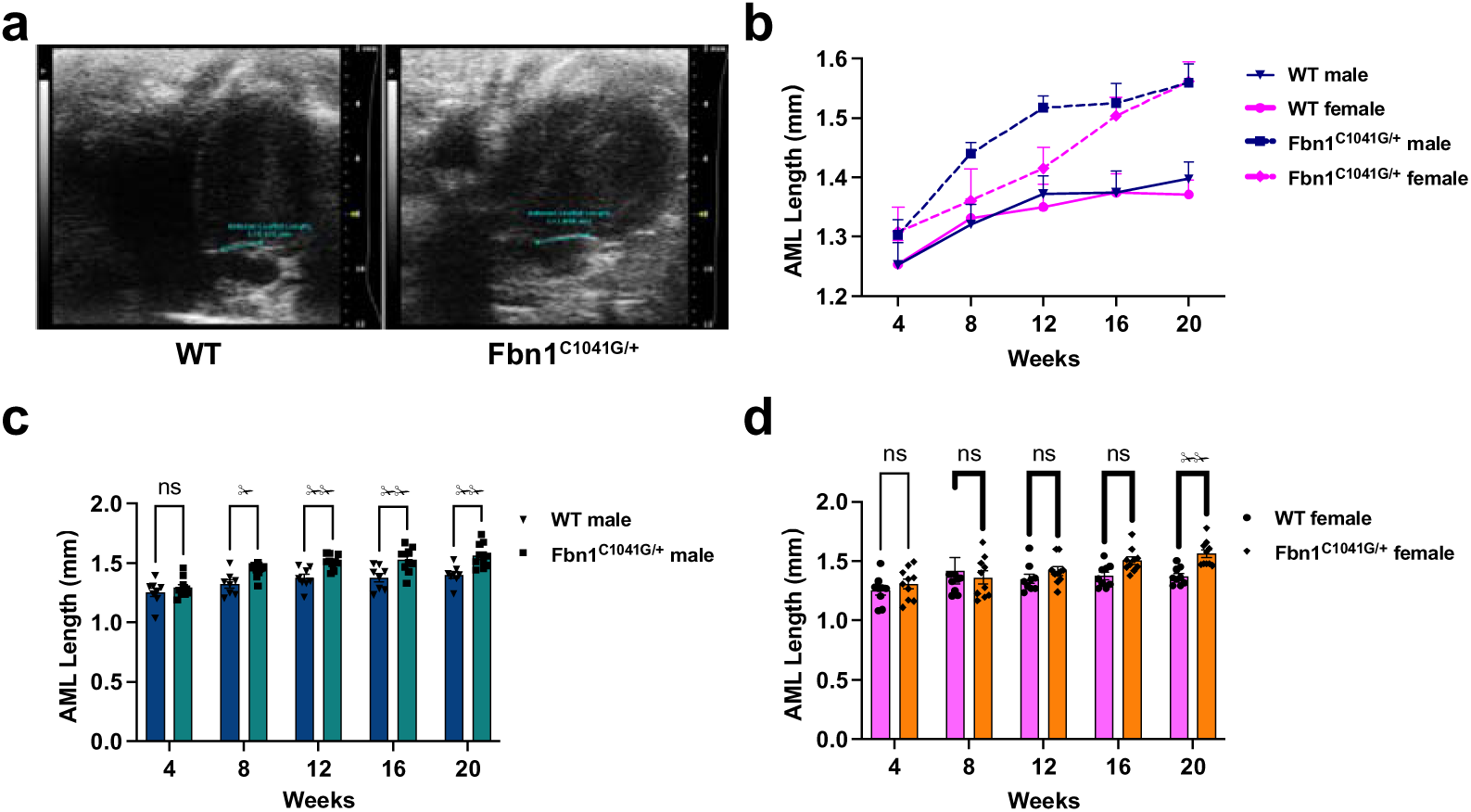
Age-related elongation of the anterior mitral leaflet in male and female *Fbn1^C1041G/+^* mice. (a) Representative transthoracic echocardiographic images showing anterior mitral leaflet length in WT and *Fbn1^C1041G/+^* mice at 20 weeks of age. (b) Age-related changes in anterior mitral leaflet length in male and female WT and *Fbn1^C1041G/+^* mice from 4 to 20 weeks of age. (c and d) Quantification of anterior mitral leaflet length in male (c) and female (d) WT and *Fbn1^C1041G/+^* mice from 4 to 20 weeks of age. Data are presented as mean ± SEM (n = 8 WT male, 9 WT female, 10 *Fbn1^C1041G/+^* male, and 10 *Fbn1^C1041G/+^*female per age group). Statistical significance was determined by two-way ANOVA followed by Sidak’s multiple comparisons test. *P < 0.05, **P < 0.01 versus age-matched WT mice; ns, not significant. AML, anterior mitral leaflet.

**Table 2.** Linear trend analysis of anterior mitral leaflet length by genotype and sex in mice aged 4 to 20 weeks.

| Group | Slope, mm/week | 95% CI | P for trend |
| --- | --- | --- | --- |
| WT male | 0.0086 | 0.0034 to 0.0138 | 0.0019 |
| WT female | 0.0070 | 0.0016 to 0.0123 | 0.0120 |
| *Fbn1*<sup>C1041G/+</sup> male | 0.0149 | 0.0105 to 0.0194 | <0.0001 |
| *Fbn1*<sup>C1041G/+</sup> female | 0.0162 | 0.0101 to 0.0223 | <0.0001 |

Note. Values represent group-specific cross-sectional linear trend analyses of anterior mitral leaflet length from 4 to 20 weeks of age. The slope represents the estimated cross-sectional difference in anterior mitral leaflet length associated with a one-week increase in age, with the 95% confidence interval (CI) shown, and is expressed in millimeters per week of age. P for trend was obtained from simple linear regression using age as a continuous variable. WT male, n = 8; WT female, n = 9; *Fbn1*<sup>C1041G/+</sup> male, n = 10; *Fbn1*<sup>C1041G/+</sup> female, n = 10 per age group.

At individual age groups, male *Fbn1^C1041G/+^* mice had longer AMLs than WT at 8, 12, 16, and 20 weeks (Fig. 2c). In females, mutant leaflet length did not differ from WT at 4-12 weeks. At 16 weeks, AML length was numerically greater in mutants than in WT, although the difference did not reach statistical significance; a significant difference emerged at 20 weeks (Fig. 2d), when mutant female values were numerically similar to those of mutant males (Fig. 2b). Given the non-significant genotype × sex × age interaction (P = 0.6744), these age-specific differences should be interpreted as cross-sectional comparisons at discrete age groups and do not provide evidence of a sex-dependent delay in leaflet elongation.

We also examined physiological sex differences in WT AML length. No significant differences were observed between male and female WT mice at any age from 4 to 20 weeks (Supplementary Fig. 3), indicating no detectable physiological sex difference in AML length under the conditions examined.

### Age- and sex-related differences in mitral inflow Doppler parameters in *Fbn1^C1041G/+^* mice

We measured mitral inflow Doppler parameters (E-wave velocity, A-wave velocity and E/A ratio) by pulsed-wave echocardiography at 4, 8, 12, 16, and 20 weeks of age (Fig. 3a). In both genotypes and sexes, E-wave and A-wave velocities showed no significant age-related changes (Fig. 3b-c and Supplementary Table 5). In contrast, the E/A ratio declined significantly with age in WT mice of either sex, whereas no significant age-related decline was observed in *Fbn1^C1041G/+^* mice of either sex (Figure 3D and Supplementary Table 5).

**Fig 3.**
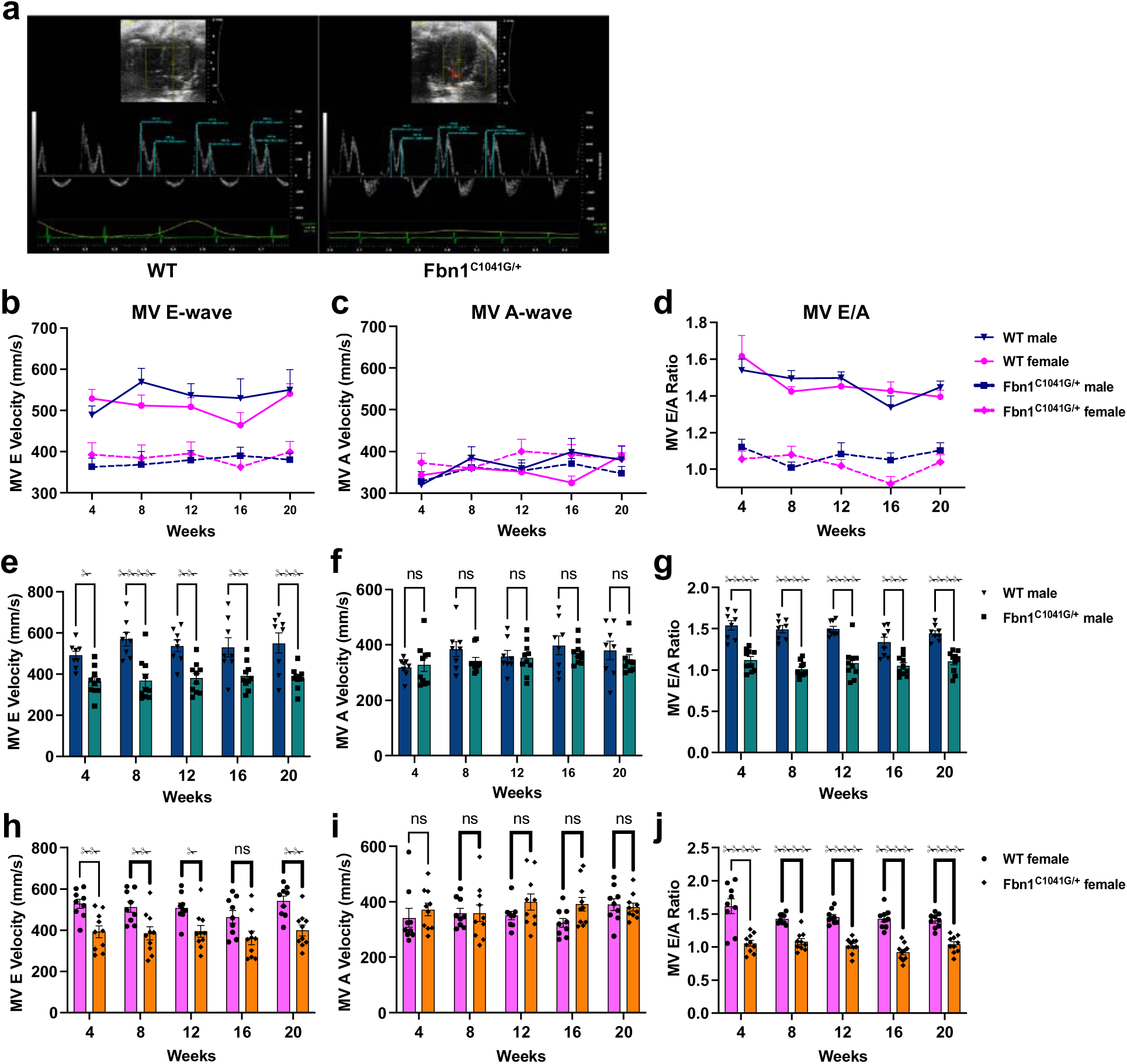
Altered mitral inflow Doppler parameters in male and female *Fbn1^C1041G/+^* mice. (a) Representative transthoracic echocardiographic pulsed-wave Doppler images of mitral inflow in WT and *Fbn1^C1041G/+^* mice at 20 weeks of age. (b-d) Age-related changes in mitral E-wave velocity, A-wave velocity, and E/A ratio in male and female WT and *Fbn1^C1041G/+^* mice from 4 to 20 weeks of age. (e-j) Quantification of mitral E-wave velocity, A-wave velocity, and E/A ratio in male (e-g) and female (h-j) WT and *Fbn1^C1041G/+^* mice. Data are presented as mean ± SEM (n = 8 WT male, 9 WT female, 10 *Fbn1^C1041G/+^* male, and 10 *Fbn1^C1041G/+^* female per age group). Statistical significance was determined by two-way ANOVA followed by Sidak’s multiple comparisons test. *P < 0.05, **P < 0.01, ***P < 0.001, ****P < 0.0001 versus age-matched WT mice; ns, not significant; MV, mitral valve.

To investigate whether these Doppler phenotypes were modulated by sex across age groups, we performed three-way ANOVA for each Doppler parameter. The genotype × sex × age interaction was not significant for E-wave velocity (P = 0.8954), A-wave velocity (P = 0.7559), or E/A ratio (P = 0.0949; Supplementary Table 4), indicating no evidence that sex-related differences in genotype effects varied across age groups. A significant genotype × sex interaction was detected for A-wave velocity (P = 0.0391), whereas the corresponding three-way interaction was not significant.

At individual age groups, male *Fbn1^C1041G/+^* mice exhibited lower E-wave velocity and E/A ratio than WT at all examined ages (Fig. 3e,g), with no genotype differences in A-wave velocity (Figure 3F). In females, E-wave velocity was lower in mutants than in WT at 4, 8, 12, and 20 weeks, with a non-significant difference at 16 weeks (Fig. 3h); A-wave velocity did not differ between genotypes at any age (Fig. 3i), whereas the E/A ratio was significantly lower in mutants at all examined ages (Fig. 3j). Thus, *Fbn1^C1041G/+^* mice of both sexes exhibited altered transmitral filling patterns characterized by lower E-wave velocity and E/A ratio with unchanged A-wave velocity. Given the non-significant genotype × sex × age interactions for all three parameters, these age-specific comparisons should not be interpreted as evidence of sex-dependent timing.

We next examined physiological sex differences in WT mice. No significant differences were observed in E-wave velocity, A-wave velocity or E/A ratio between male and female WT mice at any age (Supplementary Fig. 4a-c), indicating no detectable physiological sex difference in these transmitral filling parameters under the conditions examined.

### Age- and sex-related differences in kyphotic deformity in *Fbn1^C1041G/+^* mice

We measured kyphosis angle by lateral X-ray imaging at 4, 12 and 20 weeks of age (Fig. 4a). In *Fbn1^C1041G/+^* mice of both sexes and in WT females, the angle decreased significantly with age, with the most pronounced decline observed in mutant females; WT males showed no significant age-related change (Fig. 4b and Table 3).

**Fig 4.**
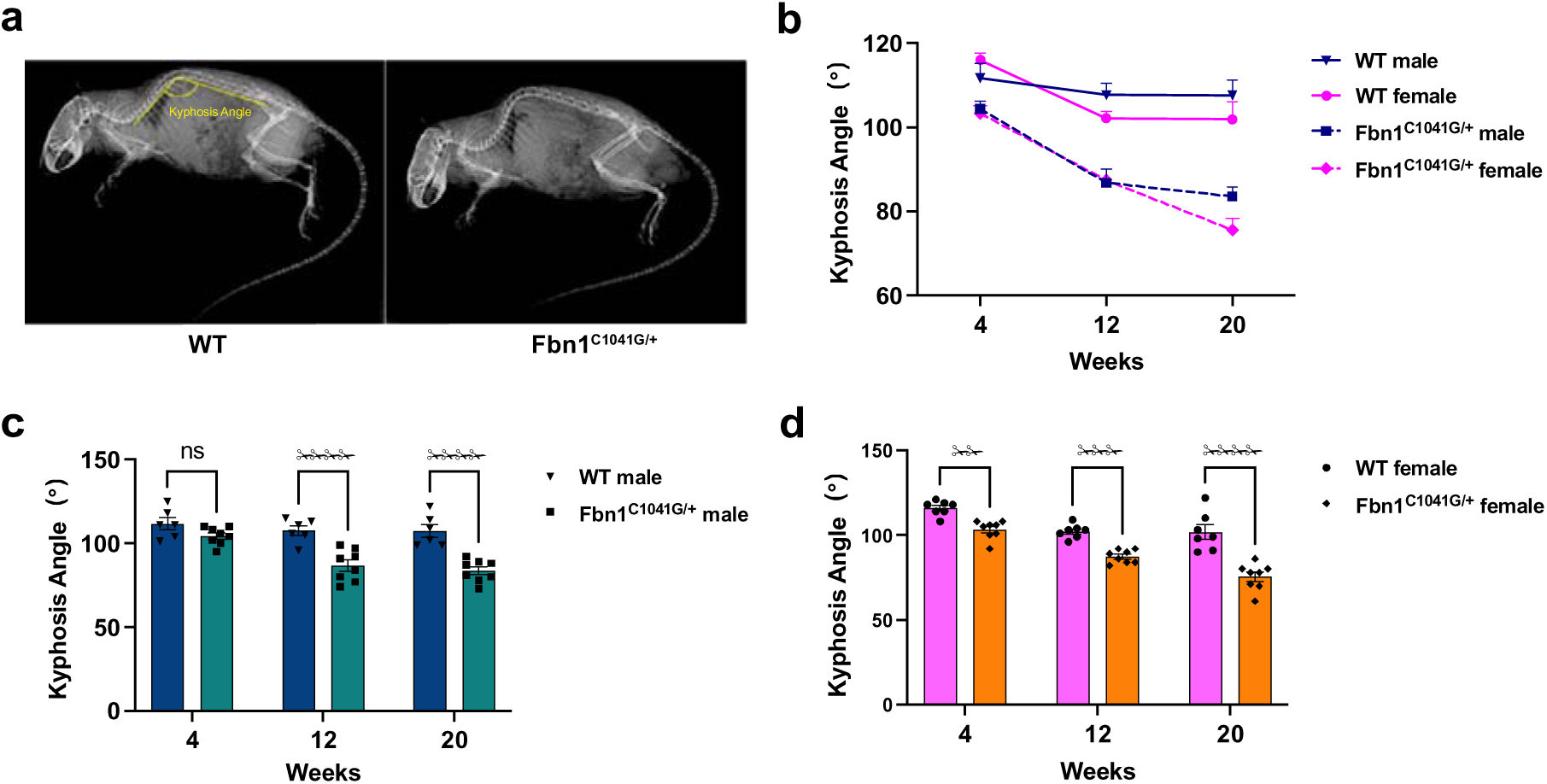
Age-related differences in kyphotic deformity in male and female *Fbn1^C1041G/+^* mice. (a) Representative lateral X-ray images showing measurement of kyphosis angle in WT and *Fbn1^C1041G/+^* mice. (b) Age-related reduction in kyphosis angle in male and female WT and *Fbn1^C1041G/+^* mice from 4 to 20 weeks of age. (c and d) Quantification of kyphosis angle in male (c) and female (d) WT and *Fbn1^C1041G/+^* mice at 4, 12, and 20 weeks of age. Data are presented as mean ± SEM (n = 8 WT male, 9 WT female, 10 *Fbn1^C1041G/+^* male, and 10 *Fbn1^C1041G/+^* female per age group). Statistical significance was determined by two-way ANOVA followed by Sidak’s multiple comparisons test. **P < 0.01, ****P < 0.0001 versus age-matched WT mice; ns, not significant.

**Table 3.**
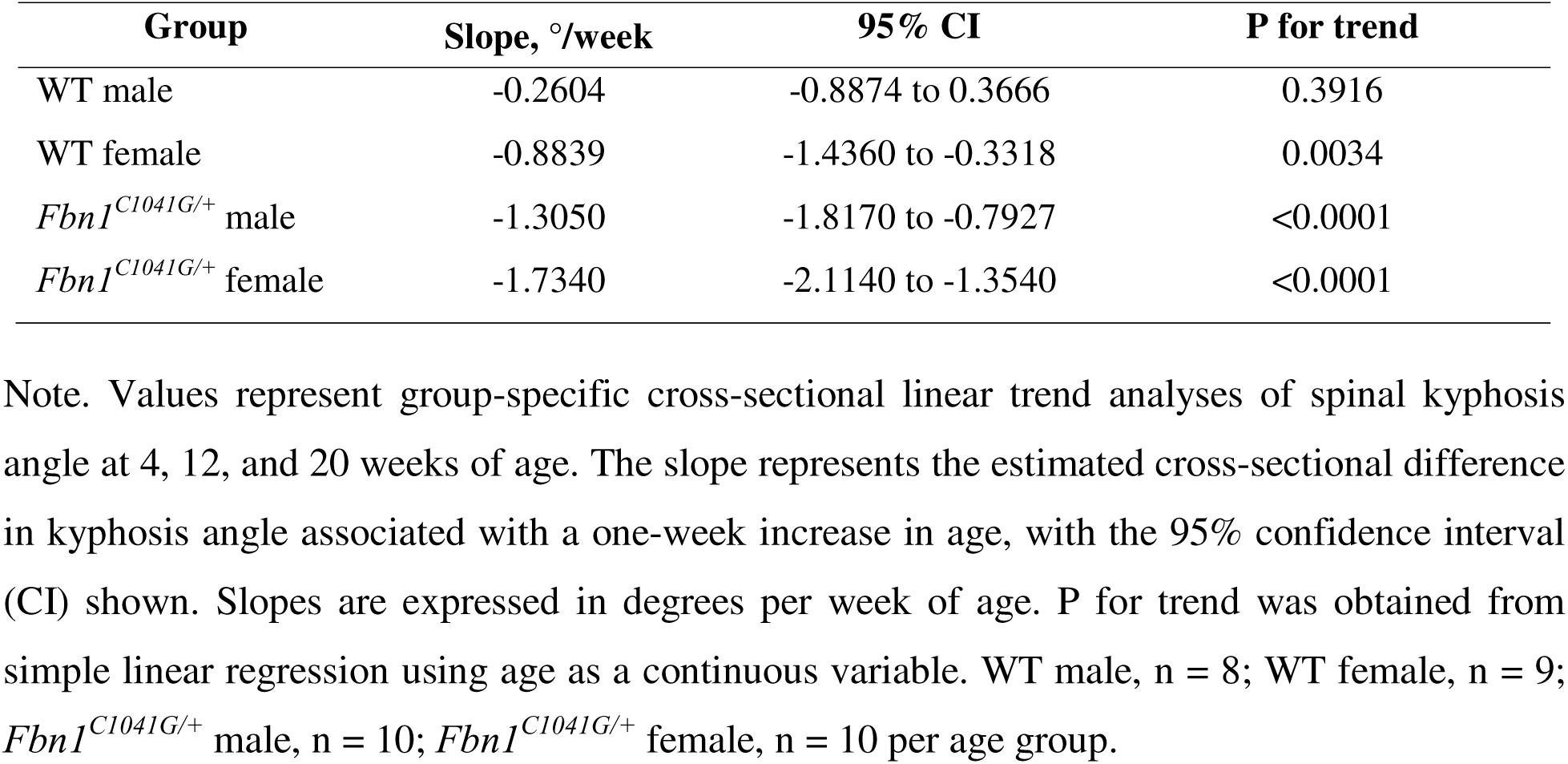
Linear trend analysis of spinal kyphosis angle by genotype and sex in mice aged 4, 12, and 20 weeks.

To formally test whether the genotype-associated kyphosis phenotype was modulated by sex, we performed three-way ANOVA. The genotype × sex × age interaction was not significant (P = 0.2915; Supplementary Table 6), indicating no evidence that the age-related genotype effect differed between males and females. Additionally, neither the genotype × sex interaction (P = 0.8582) nor the sex × age interaction (P = 0.0919) reached significance (Supplementary Table 6). Consistent with the linear trend analysis (Table 3), the genotype × age interaction was significant (P = 0.0007; Supplementary Table 6), confirming a steeper age-related decline in mutants compared with WT. WT-corrected 4-to-20-week age-group contrasts did not differ significantly between male and female mutants (Supplementary Table 7), further supporting the absence of a detectable sex-related difference in the age-associated kyphosis phenotype.

At individual age groups, male *Fbn1^C1041G/+^* mice did not differ from WT at 4 weeks but had significantly lower kyphosis angles at 12 and 20 weeks (Fig. 4c). Female mutants had significantly lower angles than WT at all three examined ages (Fig. 4d). However, given the non-significant genotype × sex × age interaction (P = 0.2915; Supplementary Table 6), these differences in the ages at which genotype comparisons reached significance should be interpreted descriptively and do not establish a sex-dependent onset of kyphotic deformity.

We also examined physiological sex differences in WT mice. Kyphosis angle did not differ significantly between male and female WT mice at any examined age (Supplementary Fig. 5), indicating no detectable physiological sex difference in spinal curvature under the conditions examined.

### Measurement reproducibility

Intraobserver and interobserver ICC point estimates ranged from 0.944 to 0.996, with CVs ranging from 0.6% to 1.7% across the echocardiographic and X-ray measurements (Supplementary Table 8).

## Discussion

Sex-related heterogeneity is increasingly recognized in connective tissue disorders [25,26], yet its influence on the early multi-system manifestations of Marfan syndrome (MFS) remains incompletely defined across age groups. In this study, we performed non-invasive phenotyping of *Fbn1^C1041G/+^* mice from 4 to 20 weeks of age to characterize age- and sex-related differences across cardiovascular and skeletal phenotypes. Distinct age-related patterns were observed across the aorta, mitral valve, transmitral filling profile, and spinal curvature, while sex-related heterogeneity was most evident in selected aortic segments. Altered transmitral filling was detectable at an earlier examined age than leaflet elongation in both sexes. Together, these observations suggest that transmitral Doppler changes may provide an early functional marker of altered filling that is detectable at earlier examined ages than overt structural leaflet elongation.

Aortic enlargement is one of the defining cardiovascular phenotypes of MFS and remains central to phenotypic assessment, surveillance, and risk stratification [1, 27]. In the present study, aortic enlargement was evident in both male and female *Fbn1^C1041G/+^* mice, but the pattern differed across aortic segments and age groups. Sex-related heterogeneity was concentrated in selected aortic segments, with the clearest differences observed in the annulus and sinus, whereas the STJ and ascending aorta showed more comparable patterns between sexes. These findings are broadly consistent with previous reports of greater susceptibility to aortic disease in male Marfan mice [7,8] and may be related to androgen-associated activation of TGF-β/ERK/SMAD signaling [28] and estrogen-associated suppression of MMP-2/MMP-9 and NF-κB signaling [29]. Importantly, female mice also developed clear aortic dilation, indicating that sex modifies the distribution of aortic involvement without conferring uniform protection. Physiological sex differences were present in several aortic dimensions in WT mice, emphasizing the importance of interpreting mutant phenotypes relative to sex-matched WT controls. Notably, the WT-corrected 4-to-20-week age-group contrasts were comparable between mutant males and females across aortic segments, suggesting that the overall magnitude of the difference between the earliest and latest examined age groups was similar between sexes. In addition, adjustment for individual body weight produced largely consistent findings, suggesting that differences in body size did not substantially account for the observed aortic enlargement.

Previous work has shown that fibrillin-1 deficiency disrupts postnatal mitral valve architecture and function, partly through enhanced TGF-β signaling and myxomatous extracellular matrix remodeling [6, 30]. In the present study, AML length increased across age groups in both genotypes, with numerically larger cross-sectional slopes in the mutant groups. Although genotype differences were detected at earlier examined ages in males than in females, the overall age-related pattern of leaflet elongation was broadly similar between sexes, providing little support for a clear sex-specific onset. The absence of physiological sex differences in WT AML length further suggests that the observed leaflet enlargement reflects the effect of the Fbn1 mutation rather than inherent differences in normal valve size. Histological reports of myxomatous changes as early as ∼1 month of age in this model [31] are compatible with our data, implying that tissue-level ECM remodeling may precede echocardiographically detectable leaflet elongation.

A notable finding was that altered transmitral filling was detectable at the earliest examined age in both sexes, whereas leaflet elongation was detected at later examined ages. Heart rate did not differ between genotypes, and no overt regurgitation was detected (data not shown), reducing the likelihood that these factors accounted for the observed Doppler differences. Similar alterations in transmitral filling have been reported in other MFS models and in patients [15,32,33]. Because ketamine/xylazine anesthesia can influence cardiac loading conditions and Doppler measurements [34], these findings should be interpreted within the context of the standardized anesthetized imaging protocol. Nevertheless, the use of identical anesthetic conditions across groups and the comparable heart rates support the validity of the between-group comparisons. Together, these observations may provide an early functional marker of altered filling that is detectable at earlier examined ages than overt structural leaflet elongation.

Kyphosis has been reported in several Fbn1-related Marfan mouse models [4, 20, 35, 36]. In the present study, a genotype-associated reduction in kyphosis angle was already detectable at 4 weeks in females, whereas it was first detected at 12 weeks in males. However, the overall age-related pattern of kyphotic deformity was broadly similar between sexes, and the WT-corrected 4-to-20-week age-group contrasts were also comparable. Thus, the difference in the earliest age at which the phenotype was detected should be interpreted as a descriptive finding rather than evidence of a sex-specific onset. Overall, kyphosis appears to be a robust skeletal manifestation of Fbn1 deficiency in both sexes.

Collectively, these findings indicate that both age and sex shape Fbn1-associated phenotypes, but their influence varies across organs and specific manifestations. Age-related differences were widespread across the cardiovascular and skeletal systems, whereas sex-related heterogeneity was concentrated mainly in selected aortic segments. These observations highlight the need to consider age and sex jointly when interpreting early Fbn1-associated phenotypes.

Several limitations should be acknowledged. First, the cross-sectional design (independent animals at each age) limits our ability to infer true within-individual longitudinal changes. Second, the sample size may have limited power to detect subtle sex-by-genotype interactions. Third, ketamine/xylazine anesthesia may have influenced transmitral Doppler measurements, although all groups were examined under the same anesthetic conditions. Finally, we did not directly examine the molecular mechanisms underlying the observed sex-related phenotypic heterogeneity. Future studies combining longitudinal phenotyping with histological, molecular, and hormone-related analyses will be important for clarifying how fibrillin-1 deficiency gives rise to age- and sex-related phenotypic heterogeneity.

Despite these limitations, our findings demonstrate that early Fbn1-associated abnormalities show distinct age-related patterns across cardiovascular and skeletal features, while sex-related differences are most evident in selected aortic segments. These findings support age-resolved and sex-stratified phenotyping in preclinical MFS studies and may help refine the selection of experimental age windows and phenotypic endpoints for mechanistic and interventional studies. The clinical relevance of these age- and sex-related patterns warrants further investigation in human MFS.

## Methods

### Mice

*Fbn1^C1041G/+^* mice, carrying a heterozygous p.C1041G mutation in *Fbn1*, were kindly provided by Dr. George Tellides (Yale University School of Medicine) [21] and maintained on a C57BL/6J background by heterozygous–WT breeding. All mice were housed under specific-pathogen-free conditions with controlled temperature (20-26°C), humidity (30-70%) and a 12:12 light-dark cycle. Animals were assigned to four groups: WT males, WT females, *Fbn1^C1041G/+^* males and *Fbn1^C1041G/+^* females. Independent cohorts were evaluated at each time point. Echocardiography was performed at 4, 8, 12, 16 and 20 weeks of age (n = 8 WT males, 9 WT females, 10 mutant males and 10 mutant females per age). X-ray imaging was performed at 4, 12 and 20 weeks using separate age-matched cohorts. Animals were allocated by genotype, sex and age. All animal experiments were approved by the Institutional Animal Care and Use Committee of Peking University People’s Hospital (Approval ID: 2022PHE027). All methods were performed in accordance with the relevant guidelines and regulations. This study is reported in accordance with the ARRIVE guidelines.

### Echocardiography

Transthoracic echocardiography was performed using a Vevo 2100 Imaging System with a 30-MHz MS400 transducer (FUJIFILM VisualSonics, Toronto, ON, Canada). Mice were anaesthetised with ketamine (80 mg/kg) and xylazine (10 mg/kg) via intraperitoneal injection and placed supine on a heated platform to maintain body temperature at 37 °C [22]. Chest hair was removed before imaging, and heart rate was monitored continuously by electrocardiography. Aortic annular, sinus of Valsalva, sinotubular junction (STJ) and ascending aortic diameters were measured at end-diastole from parasternal long-axis B-mode images, with end-diastole identified by electrocardiographic tracing [37]. Anterior mitral leaflet (AML) length was measured in the apical four-chamber view during diastole from the leaflet base to the tip [38]. Mitral inflow velocities were assessed by pulsed-wave Doppler in the same view, with the sample volume positioned at the mitral leaflet tips. Early (E-wave) and late (A-wave) diastolic velocities were averaged over three consecutive cardiac cycles, and the E/A ratio was calculated accordingly [37]. Color Doppler imaging was used to assess overt mitral regurgitation.

### X-ray imaging

For spinal curvature assessment, mice were anaesthetised as described above and placed in lateral recumbency on the detector of a digital radiography system (DR-60A(V); Shinova Medical Co., Ltd., Shanghai, China). Lateral spinal radiographs were acquired as previously described [23,24]. Kyphosis angle was measured using ImageJ software (National Institutes of Health, Bethesda, MD, USA) according to established radiographic criteria [24]. Briefly, the angle was defined by lines connecting T10 to T4 and T10 to L2 (Figure 4A); smaller angles indicated more severe kyphotic deformity.

### Blinding and measurement reproducibility

For both echocardiographic and X-ray imaging, image acquisition and quantitative measurements were performed by a genotype-blinded investigator within each imaging modality. A separate investigator who was aware of group allocation organized and verified the measurement records but did not participate in image acquisition or quantitative measurements. For reproducibility assessment, a third investigator randomly selected three mice from each genotype-sex-age stratum for each imaging parameter and anonymized the corresponding images. The original observer remeasured the anonymized images without access to the original measurements to assess intraobserver reproducibility, and a second trained observer, blinded to group allocation and the original measurements, independently measured the same images to assess interobserver reproducibility. The third investigator subsequently decoded and compiled the measurements for statistical analysis.

### Statistical analysis

Statistical analyses were performed using GraphPad Prism 10. Data are presented as mean ± SEM. Because different cohorts of animals were examined at each age, all age-related analyses reflect comparisons across independent age groups rather than longitudinal changes within individual mice, with each mouse treated as the experimental unit. Group comparisons were conducted using ordinary two-way ANOVA followed by Sidak’s multiple comparisons test, with age and genotype as factors for WT versus *Fbn1^C1041G/+^* comparisons within each sex, and age and sex as factors for physiological sex comparisons in WT mice. Linear trends across age were assessed by simple linear regression using age as a continuous variable. For aortic dimensions, additional body-weight-adjusted analyses were performed within each age-sex subgroup using multiple linear regression, with aortic diameter as the dependent variable, genotype as a categorical predictor, and body weight as a continuous covariate. For each phenotype, three-way ANOVA was performed with genotype, sex, and age as fixed factors, including all two-way interactions and the genotype × sex × age interaction. For aortic dimensions, AML length, and Doppler parameters, age was treated as a five-level categorical factor (4, 8, 12, 16 and 20 weeks); for kyphosis angle, age was treated as a three-level factor (4, 12, 20 weeks). For WT-corrected 4- to 20-week age-group contrasts, ordinary three-way ANOVA restricted to the independent 4- and 20-week cohorts was performed with age, genotype, and sex as factors, and the genotype × sex × age interaction was examined. A two-sided P < 0.05 was considered statistically significant.

For reproducibility assessment, three mice were randomly selected from each genotype-sex-age stratum for each imaging parameter, and the corresponding images were anonymized. For each imaging parameter, measurements from the selected animals were pooled across genotype, sex, and age for reproducibility analysis. Intraobserver and interobserver reproducibility were assessed using single-measure intraclass correlation coefficients (ICCs) based on a two-way random-effects model with absolute agreement. ICCs are reported with 95% confidence intervals, together with coefficients of variation (CVs). ICCs and CVs were calculated using IBM SPSS Statistics, version 27.

## Acknowledgements

The authors thank Dr. George Tellides for providing the *Fbn1^C1041G/+^* mice used in this study.

## Author Contributions

Wenxin Liu acquired and analyzed the echocardiographic data and drafted the manuscript. Shuolei Li performed X-ray imaging and anesthesia. Xiangyu Zhang collected the echocardiographic and X-ray data, verified the echocardiographic records, and performed genotyping. Lina Su performed the echocardiography. Hang Lin performed genotyping and, together with Peida Li, assisted with anesthesia and hair removal. Jingying Liu managed mouse husbandry. Yidong Niu and Sufang Li conceived, designed and supervised the study, interpreted the data, and critically revised the manuscript. All authors reviewed and approved the final version.

## Data Availability

The datasets generated and analyzed during the current study are available from the corresponding author upon reasonable request.

