## Supplementary materials for "Imaging-Based Age- and Sex-Related Cardiovascular and Skeletal Phenotypes in *Fbn1^C1041G/+^* Mice"

### Figures

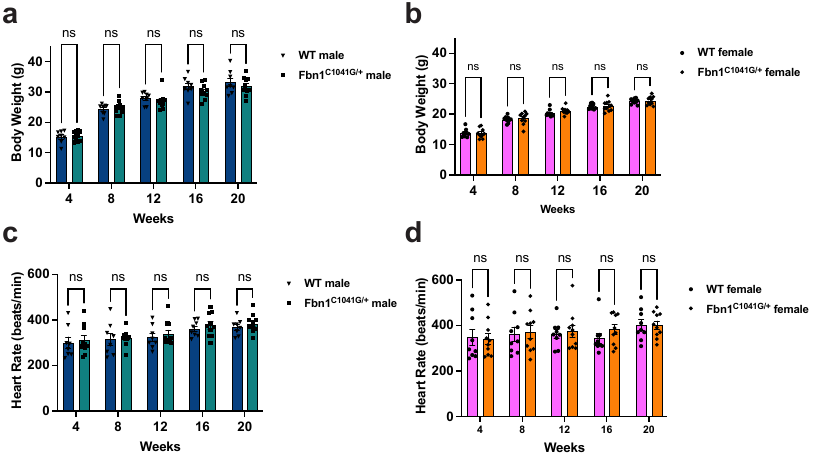

Supplementary Fig. 1. Body weight and heart rate in WT and *Fbn1^C1041G/+^* mice from 4 to 20 weeks of age. Body weight (a, b) and heart rate (c, d) measurements in male and female mice. Data are presented as mean ± SEM (n = 8 WT male, 9 WT female, 10 *Fbn1^C1041G/+^* male, and 10 *Fbn1^C1041G/+^* female per age group). Statistical significance was determined by two-way ANOVA followed by Sidak's multiple comparisons test. ns, not significant.

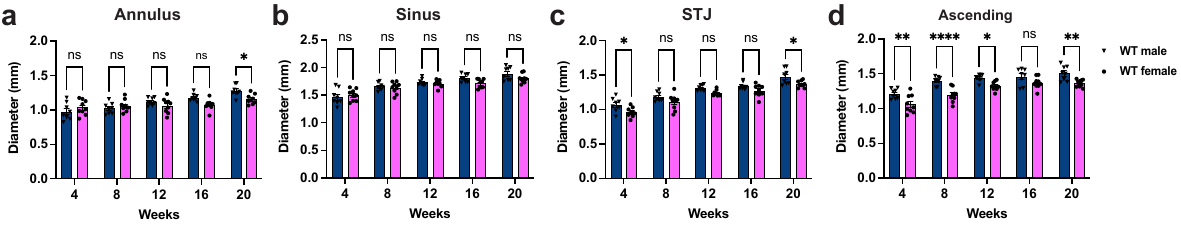

Supplementary Fig. 2. Sex-related differences in aortic root and ascending aortic diameters in WT mice. Quantification of aortic annulus (a), aortic sinus (b), sinotubular junction (c), and ascending aortic (d) diameters in male and female WT mice at 4, 8, 12, 16, and 20 weeks of age. Data are presented as mean ± SEM (n = 8 WT male and 9 WT female per age group). Statistical significance was determined by two-way ANOVA followed by Sidak's multiple comparisons test. *P < 0.05, **P < 0.01, ****P < 0.0001 between age-matched male and female WT mice; ns, not significant; STJ, sinotubular junction.

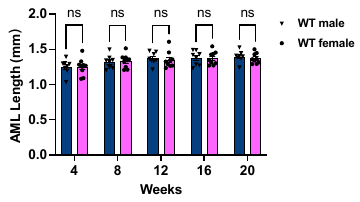

Supplementary Fig. 3. Sex-related differences in anterior mitral leaflet length in WT mice. Quantification of anterior mitral leaflet length in male and female WT mice at 4, 8, 12, 16, and 20 weeks of age. Data are presented as mean ± SEM (n = 8 WT male and 9 WT female per age group). Statistical significance was determined by two-way ANOVA followed by Sidak's multiple comparisons test. ns, not significant between age-matched male and female WT mice. AML, anterior mitral leaflet.

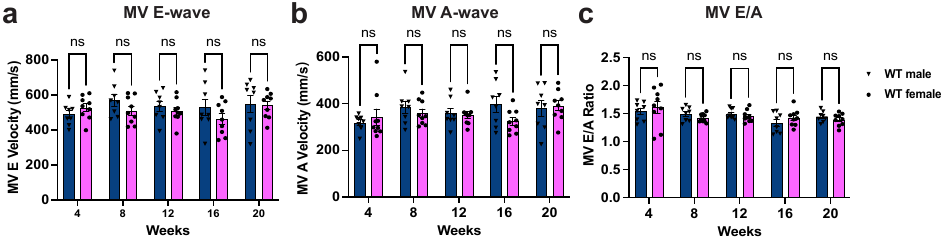

Supplementary Fig. 4. Sex-related differences in mitral inflow Doppler parameters in WT mice. (a-c) Quantification of mitral E-wave velocity (a), A-wave velocity (b), and E/A ratio (c) in male and female WT mice at 4, 8, 12, 16, and 20 weeks of age. Data are presented as mean ± SEM (n = 8 WT male and 9 WT female per age group). Statistical significance was determined by two-way ANOVA followed by Sidak's multiple comparisons test. ns, not significant between age-matched male and female WT mice. MV, mitral valve.

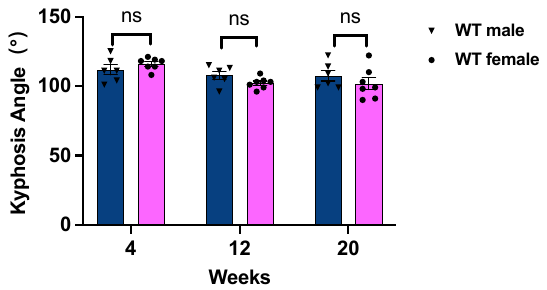

Supplementary Fig. 5. Sex-related differences in kyphosis angle in WT mice. Quantification of kyphosis angle in male and female WT mice at 4, 12, and 20 weeks of age. Data are presented as mean ± SEM (n = 8 WT male and 9 WT female per age group). Statistical significance was determined by two-way ANOVA followed by Sidak's multiple comparisons test. ns, not significant between age-matched male and female WT mice.

**Tables**

**Supplementary Table 1. Interaction statistics from three-way ANOVA of aortic dimensions by genotype, sex, and age in mice**

| **Aortic segment** | **Genotype × sex** | **Genotype × age** | **Sex × age** | **Genotype × sex × age** |
| --- | --- | --- | --- | --- |
| Annulus | F (1, 165) = 0.7566  P = 0.3857 | F (4, 165) = 4.9080  P = 0.0009 | F (4, 165) = 1.9770  P = 0.1003 | F (4, 165) = 3.7260  P = 0.0062 |
| Sinus | F (1, 165) = 15.6100  P = 0.0001 | F (4, 165) = 0.5303  P = 0.7136 | F (4, 165) = 2.0600  P = 0.0884 | F (4, 165) = 0.1986  P = 0.9388 |
| STJ | F (1, 165) = 0.7449  P = 0.3894 | F (4, 165) = 0.3293  P = 0.8580 | F (4, 165) = 0.1612  P = 0.9577 | F (4, 165) = 0.1849  P = 0.9460 |
| Ascending aorta | F (1, 165) = 0.0707  P = 0.7907 | F (4, 165) = 5.5860  P = 0.0003 | F (4, 165) = 1.1060  P = 0.3555 | F (4, 165) = 0.8911  P = 0.4707 |

Note. Values are presented as F statistics with corresponding P values from three-way ANOVA using individual-level measurements from all examined age groups (4, 8, 12, 16, and 20 weeks). Genotype (WT vs. *Fbn1^C1041G/+^*), sex (male vs. female), and age group were included as categorical factors, with all two-way interactions and the genotype × sex × age interaction included in the model. Animals examined at different ages were independent, and no repeated-measures structure was applied. The genotype × sex interaction tests whether the genotype-associated difference differs overall between males and females, whereas the genotype × sex × age interaction tests whether this sex difference in the genotype-associated effect varies across age groups. A two-sided P < 0.05 was considered statistically significant. WT male, n = 8; WT female, n = 9; *Fbn1^C1041G/+^* male, n = 10; *Fbn1^C1041G/+^* female, n = 10 per age group.

**Supplementary Table 2. Sex comparison of WT-corrected 4-to-20-week age-group contrasts in aortic dimensions in *Fbn1^C1041G/+^* mice**

| **Aortic segment** | **Sex** | **ΔWT, mm** | **ΔFbn1, mm** | **ΔΔ (WT-corrected age-group contrast), mm** | **Male − Female, mm** | **P value** |
| --- | --- | --- | --- | --- | --- | --- |
| Aortic Annulus | Male | 0.30 | 0.38 | 0.08 | -0.13 | 0.1934 |
|  | Female | 0.12 | 0.33 | **0.21** |  |  |
| Aortic Sinus | Male | 0.43 | 0.52 | 0.09 | 0.02 | 0.7133 |
|  | Female | 0.28 | 0.35 | 0.07 |  |  |
| Sinotubular Junction | Male | 0.40 | 0.43 | 0.03 | -0.02 | 0.7875 |
|  | Female | 0.41 | 0.46 | 0.05 |  |  |
| Ascending Aorta | Male | 0.30 | 0.57 | **0.27** | 0.12 | 0.1715 |
|  | Female | 0.30 | 0.45 | 0.15 |  |  |

**Note.** Values are presented as group means in millimeters. The 4-to-20-week age-group contrast (**Δ**WT or **Δ**Fbn1) was calculated as the mean diameter in the independent 20-week cohort minus that in the corresponding independent 4-week cohort. The WT-corrected age-group contrast was calculated as the 4-to-20-week contrast in *Fbn1^C1041G/+^* mice (**Δ**Fbn1) minus the corresponding contrast in WT mice (**Δ**WT) within the same sex. Male minus female represents the difference between male and female WT-corrected age-group contrasts. P values correspond to the age × genotype × sex interaction from ordinary three-way ANOVA restricted to the 4- and 20-week cohorts. This focused endpoint analysis supplements, but does not replace, the omnibus genotype × sex × age interaction analysis including all examined age groups. WT male, n = 8; WT female, n = 9; *Fbn1^C1041G/+^* male, n = 10; *Fbn1^C1041G/+^* female, n = 10 per age group.

**Supplementary Table 3. Multiple linear regression results for genotype effects on aortic dimensions after body weight adjustment**

| **Segment** | **Sex** | **Age** | **Adjusted Mean Difference (Fbn1 – WT), mm** | **95% CI** | **P value** |
| --- | --- | --- | --- | --- | --- |
| Annulus | Male | 4w | 0.1724 | 0.0730, 0.2719 | 0.0022 |
| Sinus | Male | 4w | 0.3065 | 0.1751, 0.4380 | 0.0002 |
| STJ | Male | 4w | 0.1208 | 0.0363, 0.2053 | 0.0082 |
| Ascending | Male | 4w | 0.0142 | -0.0522, 0.0806 | 0.6551 |
| Annulus | Male | 8w | 0.2780 | 0.2123, 0.3436 | <0.0001 |
| Sinus | Male | 8w | 0.3584 | 0.2177, 0.4991 | <0.0001 |
| STJ | Male | 8w | 0.1815 | 0.0895, 0.2736 | 0.0008 |
| Ascending | Male | 8w | 0.1304 | 0.0874, 0.1734 | <0.0001 |
| Annulus | Male | 12w | 0.2113 | 0.1464, 0.2762 | <0.0001 |
| Sinus | Male | 12w | 0.4353 | 0.3092, 0.5614 | <0.0001 |
| STJ | Male | 12w | 0.1774 | 0.0448, 0.3101 | 0.0121 |
| Ascending | Male | 12w | 0.1900 | 0.0707, 0.3093 | 0.0040 |
| Annulus | Male | 16w | 0.2318 | 0.1727, 0.2909 | <0.0001 |
| Sinus | Male | 16w | 0.3202 | 0.2017, 0.4387 | <0.0001 |
| STJ | Male | 16w | 0.1531 | 0.0589, 0.2474 | 0.0036 |
| Ascending | Male | 16w | 0.1830 | 0.0762, 0.2899 | 0.0025 |
| Annulus | Male | 20w | 0.2546 | 0.1719, 0.3373 | <0.0001 |
| Sinus | Male | 20w | 0.3997 | 0.2372, 0.5622 | <0.0001 |
| STJ | Male | 20w | 0.1515 | 0.0452, 0.2578 | 0.0083 |
| Ascending | Male | 20w | 0.2993 | 0.2061, 0.3924 | <0.0001 |
| Annulus | Female | 4w | 0.0546 | -0.0729, 0.1821 | 0.3775 |
| Sinus | Female | 4w | 0.1987 | 0.0475, 0.3498 | 0.0132 |
| STJ | Female | 4w | 0.1131 | 0.0182, 0.2080 | 0.0225 |
| Ascending | Female | 4w | 0.0710 | -0.0383, 0.1802 | 0.1877 |
| Annulus | Female | 8w | 0.0978 | 0.0053, 0.1903 | 0.0395 |
| Sinus | Female | 8w | 0.1938 | 0.0595, 0.3281 | 0.0075 |
| STJ | Female | 8w | 0.1308 | 0.0437, 0.2180 | 0.0058 |
| Ascending | Female | 8w | 0.1768 | 0.0808, 0.2728 | 0.0013 |
| Annulus | Female | 12w | 0.2848 | 0.2062, 0.3634 | <0.0001 |
| Sinus | Female | 12w | 0.2047 | 0.1123, 0.2970 | 0.0002 |
| STJ | Female | 12w | 0.1191 | 0.0720, 0.1662 | <0.0001 |
| Ascending | Female | 12w | 0.1732 | 0.0745, 0.2719 | 0.0019 |
| Annulus | Female | 16w | 0.3355 | 0.2351, 0.4360 | <0.0001 |
| Sinus | Female | 16w | 0.2133 | 0.0862, 0.3403 | 0.0026 |
| STJ | Female | 16w | 0.1134 | 0.0222, 0.2046 | 0.0180 |
| Ascending | Female | 16w | 0.2075 | 0.0916, 0.3233 | 0.0016 |
| Annulus | Female | 20w | 0.2720 | 0.1702, 0.3739 | <0.0001 |
| Sinus | Female | 20w | 0.2527 | 0.1641, 0.3412 | <0.0001 |
| STJ | Female | 20w | 0.1644 | 0.0896, 0.2392 | 0.0003 |
| Ascending | Female | 20w | 0.2112 | 0.1049, 0.3176 | 0.0007 |

Note. Adjusted mean differences (Fbn1 − WT) and 95% confidence intervals were estimated by multiple linear regression within each age-sex subgroup, with genotype included as a categorical predictor and body weight as a continuous covariate. Positive values indicate larger aortic diameters in *Fbn1^C1041G/+^* mice than in WT controls. Animals examined at different ages were independent, and no repeated-measures structure was applied. A two-sided P < 0.05 was considered statistically significant. WT male, n = 8; WT female, n = 9; *Fbn1^C1041G/+^* male, n = 10; *Fbn1^C1041G/+^* female, n = 10 per age group.

**Supplementary Table 4. Interaction statistics from three-way ANOVA of anterior mitral leaflet length and Doppler parameters by genotype, sex, and age in WT and *Fbn1^C1041G/+^*** **mice**

| **Phenotype** | **Genotype × sex** | **Genotype × age** | **Sex × age** | **Genotype × sex × age** |
| --- | --- | --- | --- | --- |
| AML length | F (1, 165) = 1.0790  P = 0.3005 | F (4, 165) = 2.1310  P = 0.0791 | F (4, 165) = 0.5563  P = 0.6947 | F (4, 165) = 0.5844  P = 0.6744 |
| E-wave velocity | F (1, 165) = 1.8130  P = 0.1800 | F (4, 165) = 0.3932  P = 0.8133 | F (4, 165) = 1.0780  P = 0.3693 | F (4, 165) = 0.2725  P = 0.8954 |
| A-wave velocity | F (1, 165) = 4.324  P = 0.0391 | F (4, 165) = 0.7503  P = 0.5592 | F (4, 165) = 1.2160  P = 0.3058 | F (4, 165) = 0.4725  P = 0.7559 |
| E/A ratio | F (1, 165) = 1.3620  P = 0.2449 | F (4, 165) = 1.089  P = 0.3637 | F (4, 165) = 0.3784  P = 0.8238 | F (4, 165) = 2.0140  P = 0.0949 |

Note. Values are presented as F statistics with corresponding P values from ordinary three-way ANOVA using individual-level measurements from all examined age groups (4, 8, 12, 16, and 20 weeks). Genotype (WT vs. *Fbn1^C1041G/+^*), sex (male vs. female), and age group were included as categorical factors, with all two-way interactions and the genotype × sex × age interaction included in the model. Animals examined at different ages were independent, and no repeated-measures structure was applied. A two-sided P < 0.05 was considered statistically significant. **Doppler parameters** include E-wave velocity, A-wave velocity and E/A ratio. WT male, n = 8; WT female, n = 9; *Fbn1^C1041G/+^* male, n = 10; *Fbn1^C1041G/+^* female, n = 10 per age group. AML, anterior mitral leaflet.

**Supplementary Table 5. Linear trend analysis of mitral inflow Doppler parameters by genotype and sex in mice aged 4 to 20 weeks**

| **Mitral inflow parameter** | **Group** | **Slope** | **95% CI** | **P for trend** |
| --- | --- | --- | --- | --- |
| E-wave velocity | WT male | 2.0330 | -3.9010 to 7.9670 | 0.4921 |
|  | WT female | -0.5855 | -4.7350 to 3.5640 | 0.7774 |
|  | *Fbn1^C1041G/+^* male | 1.4130 | -2.1990 to 5.0260 | 0.4354 |
|  | *Fbn1^C1041G/+^* female | -0.2325 | -4.8110 to 4.3460 | 0.9191 |
| A-wave velocity | WT male | 3.3830 | -0.8412 to 7.6070 | 0.1132 |
|  | WT female | 1.5000 | -2.0720 to 5.0720 | 0.4017 |
|  | *Fbn1^C1041G/+^* male | 1.1990 | -1.8360 to 4.2330 | 0.4310 |
|  | *Fbn1^C1041G/+^* female | 1.2910 | -2.6430 to 5.2250 | 0.5126 |
| E/A ratio | WT male | -0.0086 | -0.0166 to -0.0007 | 0.0342 |
|  | WT female | -0.0110 | -0.0207 to -0.0013 | 0.0277 |
|  | *Fbn1^C1041G/+^* male | 0.0002 | -0.0068 to 0.0071 | 0.9654 |
|  | *Fbn1^C1041G/+^* female | -0.0048 | -0.0116 to 0.0020 | 0.1595 |

**Note.** Values represent group-specific cross-sectional linear trend analyses of mitral inflow Doppler parameters from 4 to 20 weeks of age. The slope represents the estimated cross-sectional difference in the corresponding parameter associated with a one-week increase in age, with the 95% confidence intervals (CIs) shown. P for trend was obtained from simple linear regression using age as a continuous variable. For E-wave and A-wave velocities, slopes are expressed as mm/s per week of age; for the E/A ratio, slopes are expressed as ratio units per week of age. WT male, n = 8; WT female, n = 9; *Fbn1^C1041G/+^* male, n = 10; *Fbn1^C1041G/+^* female, n = 10 per age group.

**Supplementary Table 6. Interaction statistics from three-way ANOVA of kyphosis angle by genotype, sex, and age in WT and *Fbn1^C1041G/+^*** **mice**

| **Phenotype** | **Genotype × sex** | **Genotype × age** | **Sex × age** | **Genotype × sex × age** |
| --- | --- | --- | --- | --- |
| Kyphosis angle | F (1, 75) = 0.0321  P = 0.8582 | F (2, 75) = 7.9760  P = 0.0007 | F (2, 75) = 2.4650  P = 0.0919 | F (2, 75) = 1.2530  P = 0.2915 |

Note. Values are presented as F statistics with corresponding P values from ordinary three-way ANOVA using individual-level measurements from all examined age groups (4, 12, and 20 weeks). Animals examined at different ages were independent, and no repeated-measures structure was applied. A two-sided P < 0.05 was considered statistically significant. WT male, n = 8; WT female, n = 9; *Fbn1^C1041G/+^* male, n = 10; *Fbn1^C1041G/+^* female, n = 10 per age group.

**Supplementary Table 7. Sex comparison of WT-corrected 4-to-20-week age-group contrasts in kyphosis angle in *Fbn1^C1041G/+^* mice**

| **Sex** | **ΔWT, °** | **ΔFbn1, °** | **ΔΔ (WT-corrected age-group contrast), °** | **Male − Female, °** | | **P value** |
| --- | --- | --- | --- | --- | --- | --- |
| Male | -4.17 | -20.88 | -16.71 | -3.10 | | 0.6972 |
| Female | -14.14 | -27.75 | -13.61 | |  |  |

**Note.** Values are presented as group means in degrees. The 4-to-20-week age-group contrast (ΔWT or ΔFbn1) was calculated as the mean kyphosis angle in the independent 20-week cohort minus that in the corresponding independent 4-week cohort. The WT-corrected age-group contrast was calculated as the 4-to-20-week contrast in *Fbn1^C1041G/+^* mice (ΔFbn1) minus the corresponding contrast in WT mice (ΔWT) within the same sex. Male minus female represents the difference between male and female WT-corrected age-group contrasts. P value corresponds to the age × genotype × sex interaction from ordinary three-way ANOVA restricted to the 4- and 20-week cohorts. This focused endpoint analysis supplements, but does not replace, the omnibus genotype × sex × age interaction analysis including all examined age groups. For kyphosis angle, a negative contrast indicates a smaller angle and therefore more severe kyphosis. WT male, n = 8; WT female, n = 9; *Fbn1^C1041G/+^* male, n = 10; *Fbn1^C1041G/+^* female, n = 10 per age group.

**Supplementary Table 8. Intraobserver and interobserver reproducibility of echocardiographic and kyphosis measurements**

| **Parameter** | **Intraobserver ICC (95% CI)** | **Intraobserver CV (%)** | **Interobserver ICC (95% CI)** | **Interobserver CV (%)** |
| --- | --- | --- | --- | --- |
| Aortic annulus | 0.985 (0.975-0.991) | 1.1 | 0.991 (0.985-0.995) | 0.8 |
| Aortic sinus | 0.989 (0.982-0.993) | 0.9 | 0.994 (0.990-0.996) | 0.6 |
| STJ | 0.981 (0.960-0.990) | 1.2 | 0.988 (0.972-0.994) | 0.9 |
| Ascending aorta | 0.985 (0.974-0.991) | 0.8 | 0.988 (0.979-0.993) | 0.7 |
| AML length | 0.957 (0.894-0.979) | 1.5 | 0.944 (0.890-0.970) | 1.7 |
| E-wave velocity | 0.994 (0.988-0.997) | 1.0 | 0.996 (0.992-0.998) | 1.0 |
| A-wave velocity | 0.992 (0.986-0.995) | 0.9 | 0.995 (0.991-0.997) | 0.9 |
| E/A ratio | 0.984 (0.973-0.990) | 1.5 | 0.990 (0.983-0.994) | 1.3 |
| Kyphosis angle | 0.989 (0.978-0.994) | 1.0 | 0.993 (0.987-0.997) | 0.8 |

**Note.** Values are presented as intraclass correlation coefficients (ICCs) with 95% confidence intervals (CIs) and coefficients of variation (CVs). Intraobserver reproducibility was assessed using two repeated measurements obtained by the same observer. Interobserver reproducibility was assessed by comparing the original measurement obtained by the first observer with an independent measurement obtained by the second observer. For each imaging parameter, measurements were pooled across genotype, sex, and age for reproducibility analysis. The pooled sample comprised 60 mice for each echocardiographic parameter and 36 mice for kyphosis angle. ICCs were calculated using a two-way random-effects model with absolute agreement and are reported as single-measure estimates. CVs are expressed as percentages. AML, anterior mitral leaflet; STJ, sinotubular junction.
