## Supplementary figures and images for "Imaging-Based Age- and Sex-Related Cardiovascular and Skeletal Phenotypes in *Fbn1^C1041G/+^* Mice"

### Supplementary Figure 1

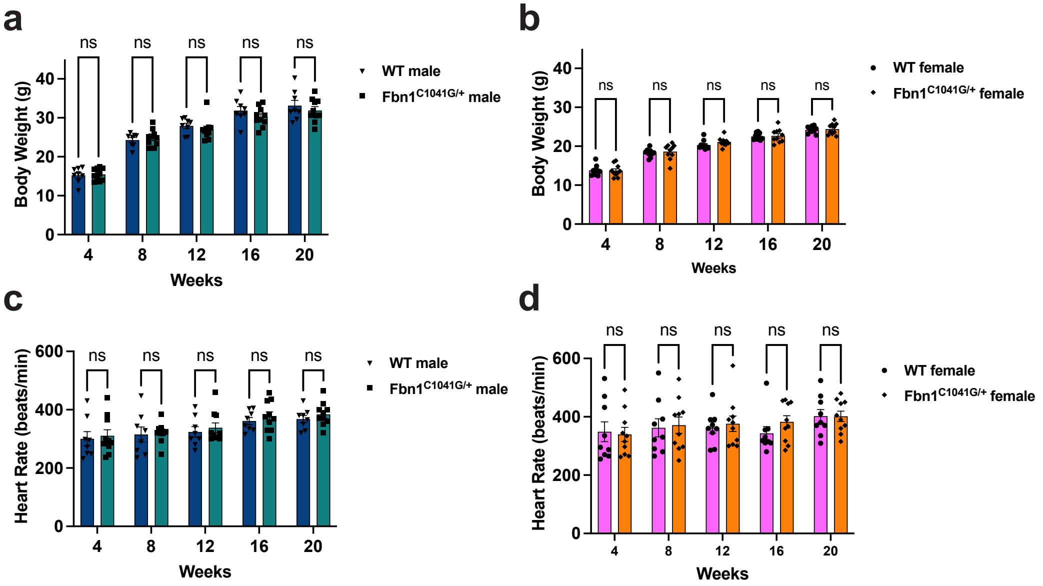

### Supplementary Figure 2

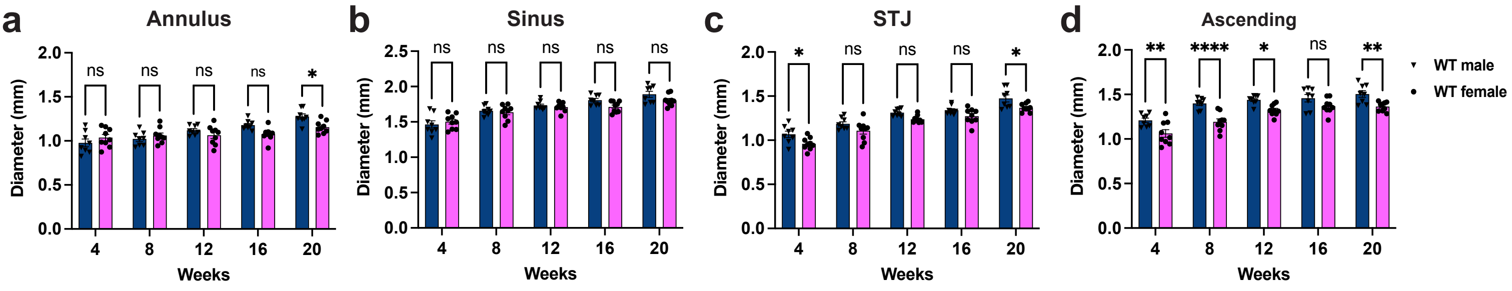

### Supplementary Figure 3

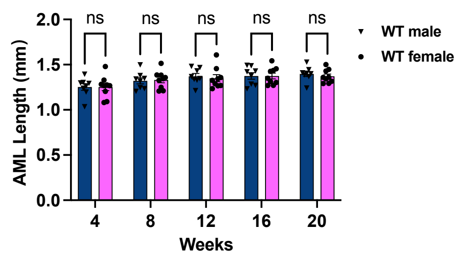

### Supplementary Figure 4

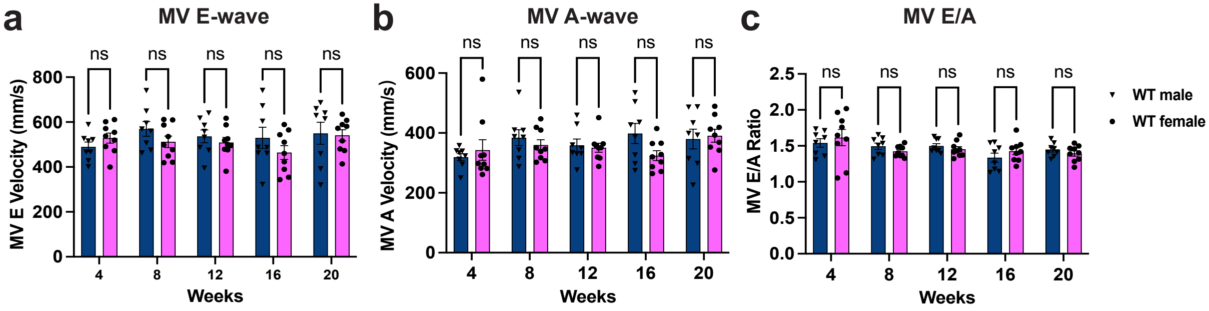

### Supplementary Figure 5

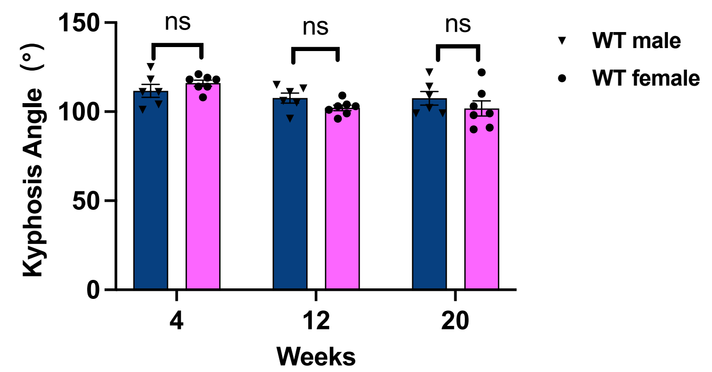
